# *Ex Vivo* Testing of Inflatable Penile Prosthesis in Human Cadaveric Penis with Paired *in Silico* Model offering Surgical and Biomechanical Insights

**DOI:** 10.1101/2025.11.20.689496

**Authors:** Majid Akbarzadeh Khorshidi, Shirsha Bose, Ivor M. Cullen, John Sullivan, Robert Johnston, Kenneth Patterson, Brian Watschke, Thomas Sinnott, Evania Mareena, Caitríona Lally

**Author notes:** Corresponding author: Caitriona Lally. Both the authors are joint-first authors on this paper (Majid Akbarzadeh Khorshidi, Shirsha Bose).

## Abstract

**Background:** Inflatable penile prostheses (IPPs) are a critical solution for patients with erectile dysfunction refractory to medical therapy. However, a detailed understanding of their mechanical interaction with penile tissues remains limited.

**Aim:** To develop and validate an innovative experimental-computational framework for studying IPP behaviour through ex vivo implantation and inflation testing in human cadaveric penile tissue, paired with a finite element-based (FE-based) computational model.

**Methods:** An AMS 700 IPP was surgically implanted into a human cadaveric penis (including the glans and ∼15 cm of shaft) using standard clinical techniques. The cylinders were placed within the corpora cavernosa, with the pump and reservoir positioned externally in a closed hydraulic loop. An inflation test was performed ex vivo, with real-time ultrasound imaging used to monitor cylinder expansion. Internal pressure was recorded using a digital barometer. Following inflation, sectional analysis enabled 3D approximation of penile shaft geometry. A representative FE-based computational model was developed, incorporating anatomically accurate tissue layers—tunica albuginea (TA), corpus cavernosa (CC), corpus spongiosum (CS), and fascia—with realistic material properties to simulate the inflation process.

**Outcomes:** This study enabled the mechanical response of penile tissues to IPP inflation to be quantified using both experimental and computational modalities.

**Results:** The combined use of ultrasound imaging and digital pressure monitoring successfully captured dynamic IPP behaviour during inflation. The FE model reproduced experimental outcomes with good fidelity, providing a detailed understanding of stress distribution and tissue deformation.

**Clinical Implications:** This integrated approach can inform future IPP design improvements and aid surgeons in preoperative planning by offering predictive insights into prosthesis–tissue interaction.

**Strengths & Limitations:** A major strength of this study is the novel integration of cadaveric experimentation with computational modelling. However, limitations include the absence of active physiological responses and potential variability due to cadaveric tissue properties.

**Conclusion:** This pioneering work establishes a robust platform for studying IPP mechanics in realistic anatomical contexts, with promising implications for optimising device design and improving patient outcomes in urologic surgery.

## 1. Introduction

Erectile dysfunction (ED) is the inability to achieve or maintain an erection in the male reproductive organ and affects approximately 52% of males over 40 [1]–[3]. Implantation of a penile prosthesis is the most common treatment for patients who do not respond well to pharmacological treatments [4]. Various types of penile prostheses are being used to resolve ED [2], [5], [6]. Two general types of penile prostheses are malleable penile prostheses (MPPs), where the girth and stiffness always remain the same [5] and inflatable penile prostheses (IPPs) [2]. IPP is the most advanced device for ED treatment with a high rate of satisfaction [7], whereby the IPP can be controlled in terms of girth, length, and stiffness. The three piece IPP, such as the AMS 700, the Coloplast Titan, and the Rigicon Infla 10 are the most popular penile prostheses [2], [8] and consist of two cylinders implanted in the corpus cavernosal cavities of the penis, a reservoir of saline placed in the space of retzius or ectopically in the abdominal wall musculature in the lower abdomen, and a mechanical pump placed inside the scrotum. These are connected to each other by resilient tubes as shown in Fig. 1a. Squeezing the pump moves fluid into the cylinders, creating rigidity in the prosthesis and thereby an erection. Releasing the device by pressing the deflate button on the pump returns the penis to a flaccid state. AMS 700 CX only expands radially while the length remains constant, while AMS 700 LGX expands in both length and girth.

**Fig. 1.**
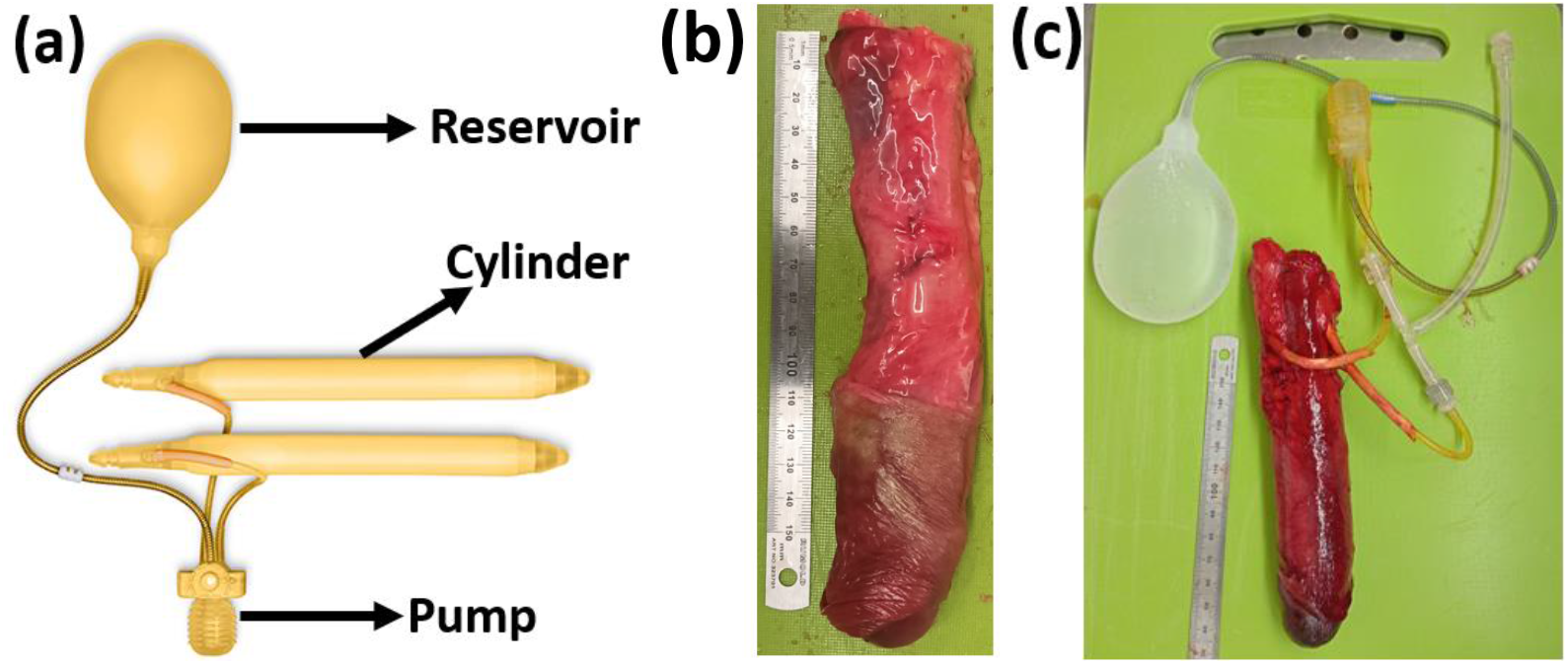
(a) Boston Scientific AMS 700 CX IPP device [27], (b) human cadaveric penis, and (c) IPP device implanted into the penis.

The early IPPs were developed in 1970s, however, there has been limited research on understanding the effect of inflation-deflation of the device on the neighbouring tissues [2]. Some post-IPP implantation complications have been reported based on computed tomography (CT) scans, such as severe tissue swelling and thickening [9]. Other complications include tissue erosion, malposition/migration, IPP mechanical failure, and hematomas which can severely affect the patient’s life post-IPP implantation [9]–[12]. Although device related complications are reported in the literature, there is still a lack of appropriate in vitro or in silico testbeds which would assist in addressing these issues.

It is worth mentioning that designing an optimal medical device with good long-term performance requires preclinical testing in both *ex-vivo* and *in silico* environments, as conducting *in vivo* tests on new invasive devices is limited to regulatory approved clinical trials (Regulation (EU) 2017/745 on medical devices (MDR) and Regulation (EU) 2017/746 (IVDR) on in vitro diagnostic medical devices). Few studies have performed *ex vivo* IPP implantation on cadaveric penis [13], [14], or *ex vivo* mechanical tests on IPP devices [15], [16]. The challenges associated with acquiring and testing human tissue, including cost and ethical concerns [17], [18], make human tissue experimentation difficult and expensive. On the other hand, due to the fact that animal penile tissues differ from human penile tissues in both mechanical properties [19] and anatomy [2], getting experimental data from animal tissues is not the ideal choice and involves significant discrepancies [2], [19]. Therefore, the development of preclinical models is crucial for advancing the study of human penile tissue mechanics. Preclinical models seek to evaluate penile prostheses’ performance and behaviour within penile tissues [20], improving understanding of post-implantation complications and long-term issues [10], and for surgical training and simulator purpose [14], [21].

Computational modelling can strongly contribute to enhance the understanding of penile tissue mechanics, predict IPP performance, and optimise design. Finite element (FE) analysis has been used to simulate the mechanical interactions between the prosthesis and penile tissues[20], [22], [23], many limited to simple models which did not account for the penile tissue’s non-linear behaviour or anisotropy [19], [21], [22]. Such models will therefore not be able to predict the tissue’s deformation precisely during medical device implantation. Several computational models have been developed to study the deformation, anisotropy and stress distributions in penile tissues [19], [20], [23]–[25]. However, those models were not informed by mechanically tested human tissue data. Previous work from our group, presented in Akbarzadeh Khorshidi et al. [26], utilised an inverse FE approach to characterise human penile tissue, leading to more realistic predictions for the tissues’ mechanical responses. To-date very limited research has been performed on IPPs using computational models based on experimental data from human penile tissues.

The objective of this study is to establish a preclinical testing approach by performing *ex vivo* IPP implantation and inflation test into human cadaveric penis combined with an accurate FE model. The FE model incorporated experimentally derived human penile tissue properties [26] together with implant specifications from the project partner (Boston Scientific) to enable prediction of tissue stress and strain distributions during IPP inflation. This *in silico* model offers a preclinical testbed to understand the IPP device performance and perform mechanical analysis.

## 2. Methods

### 2.1. Ex vivo IPP implantation

A fresh human cadaveric penis with a post-mortem time interval of 1–5 days from a donor aged 85 years of age received from ScienceCare, USA. The tissue was received in a frozen state and stored at -20 °C. After defrosting at room temperature, the penis was cleaned by removing unwanted tissues in such a way that foreskin, glans and 15 cm penile shaft remained, as shown in Fig. 1b.

Boston Scientific AMS 700™ CX with cylinders of 12 mm initial diameter and 15 cm total length, a 100 ml reservoir, and MS (Momentary Squeeze) Pump, was used to be implanted into the cadaveric tissue for IPP inflation test. An incision was made in the proximal penile cadaveric shaft skin ventrally. A standard 3cm corporotomy was made in the proximal tunica albuginea and a single pass furlow dilatation was performed in the corporal smooth muscle to allow implantation of a 15cm IPP in both corpora. The corporotomies were closed in a standard fashion around the tubing using 0 Vicry suture.

### 2.2. IPP inflation test

The inflation test was conducted using the human cadaveric penis and Boston Scientific’s AMS 700 CX IPP device (detailed in section 2.1). A digital barometer was attached into the *ex vivo* test set-up to measure the inflation pressure during the test. The cylinders inflation within the tissue was visualised using an ultrasound scanner -SIEMENS ACUSON S2000™-equipped with a 9L4 linear probe operating at 8 MHz frequency. An image was taken at each pressure increment from 0 to 20 psi and at five different locations through the penis length (every 20 mm from the distal region to the proximal region, covering 80 mm length of the cadaveric penis, see Fig. 4). The IPP inflation test was done in a water bath (PBS, 37°C), while the cadaveric penis and the IPP cylinders implanted were placed inside the bath and the barometer, pump, and reservoir remained outside. The IPP cylinders within the penis were pressurised by manually compressing the pump, while the ultrasound images were captured at each pressure increment for subsequent postprocessing to measure the IPP cylinder diameters (see Fig. 3).

### 2.3. Finite element modelling

This geometry was generated by creating cross-sectional sketches based on photographs of five distinct sections from the cadaveric penis. These cross sections were obtained by cutting the penile shaft at 20 mm intervals along its length, from the distal to the proximal end (after the IPP inflation test). The five sketches were lofted in Abaqus software (Simulia 2022) to form a 3D model of the penis, as shown in Fig. 2. A 12 mm diameter hole was added to the CC layer to accommodate the IPP cylinders. A pressure of 137 kPa (≈20 psi) was applied to the internal surface of the IPP cylinder, based on specifications from Boston Scientific internal documents, which indicate 20 psi as the maximum operating pressure. Both ends of the penile shaft were restricted to move in the longitudinal direction.

**Fig. 2.**
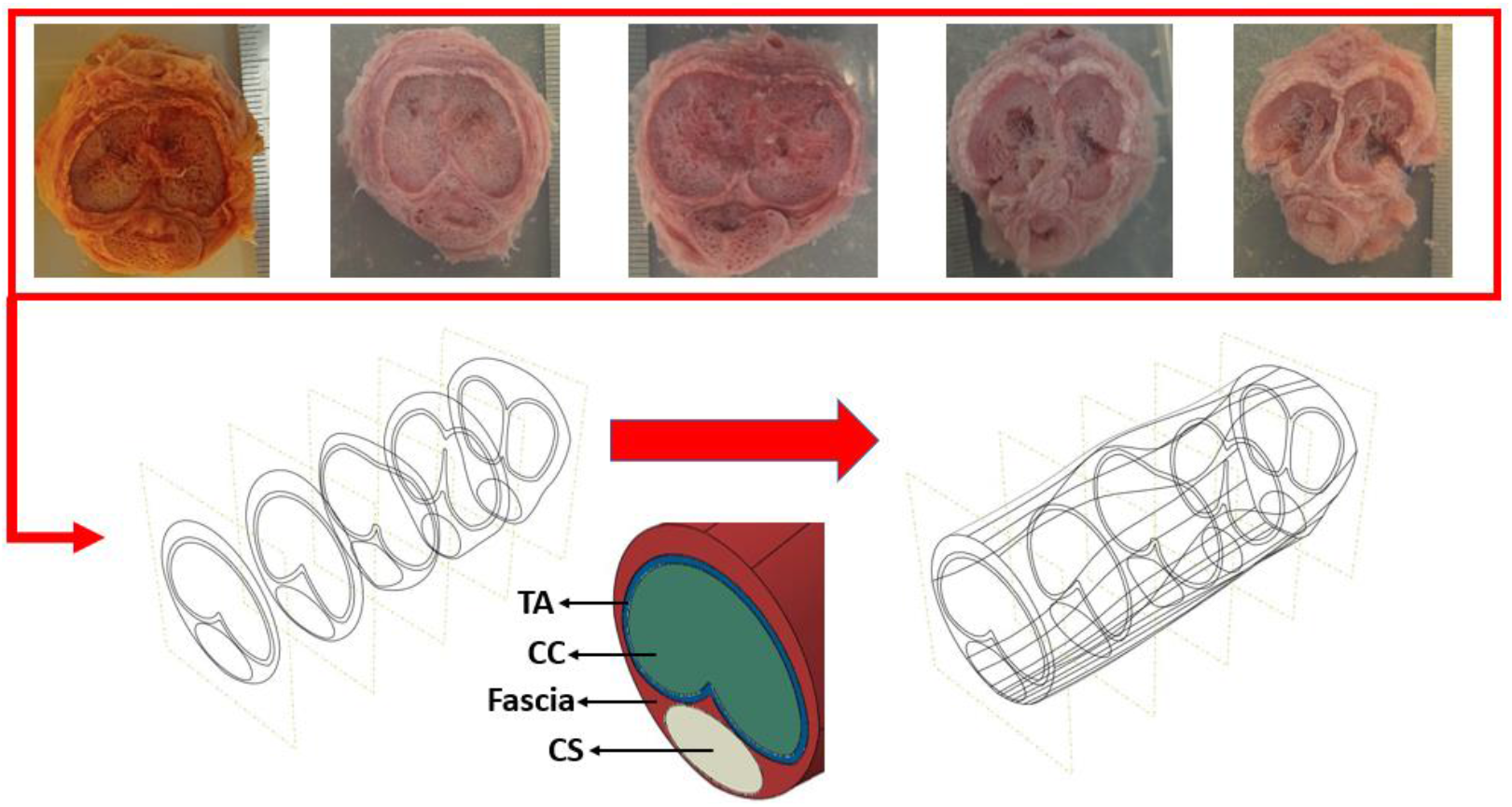
3D geometry of penis including four individual layers: TA, CC, CS, and fascia.

The penile tissue was defined to consist of four individual layers: TA, CC, CS, and fascia layer (see Fig. 2). Hyperelastic material models were employed to define the properties of each tissue layer: the Holzapfel-Gasser-Ogden anisotropic model for TA (with circumferential collagen fibres orientation), the Ogden model for CS, CC, and IPP cylinders, and the neo-Hookean model for the fascia layer. The tissue parameters used were adopted from our previous research on human penile tissue characterisation [26], see Table 1. The material properties of the IPP cylinder were obtained through an inverse FE simulation to replicate the unconstrained pressure-diameter experimental data provided in Boston Scientific documents [28].

**Table 1:** Human penile tissue properties [26] and IPP cylinder properties [28].

| Tissue Layer | Material Parameters |  |  |  |  |  |
| --- | --- | --- | --- | --- | --- | --- |
| TA (HGO model) | $C_{10}=100$ kPa | $k_1=1500$ kPa | $k_2=0.1$ | $\kappa=0$ | $D=1e-5$ kPa$^{-1}$ | |
| CC (Ogden model) | $\mu=0.022$ kPa | $\alpha=20$ | | | $D=1.8$ kPa$^{-1}$ | |
| CS (Ogden model) | $\mu_1=0.6$ kPa | $\alpha=6$ | | | $D=0.1$ kPa$^{-1}$ | |
| Fascia (neo-Hookean model) | $C_{10}=0.025$ kPa | $D=0.8$ kPa$^{-1}$ | | | | |
| IPP cylinder (Ogden model) | $\mu_1=0.85$ kPa | $\mu_2=629.76$ kPa | $\alpha_1=19.43$ | $\alpha_2=0.21$ | $D_1=5e-5$ kPa$^{-1}$ | $D_2=1e-5$ kPa$^{-1}$ |

## 3. Results

The post-processing of ultrasound images enabled measurement of the IPP cylinder diameter at various inflation pressures and locations, as shown in Fig. 3. Using these measurements, the *ex vivo* experimental data are plotted to generate a diameter-pressure diagram of the IPP device, illustrating its inflation behaviour within cadaveric tissue (see Fig. 4). The FE simulation results of IPP inflation are then compared with the experimental data, as demonstrated in Fig. 4. The comparison indicates that the *in silico* FE-based model accurately replicates the *ex vivo* IPP inflation test. The *ex vivo* pressure-inflation data reveals the nonlinear behaviour of the IPP cylinder within the tissue, which is consistent with the nonlinear inflation response observed when the cylinder is inflated without tissue [28], see Fig. 4.

**Fig. 3.**
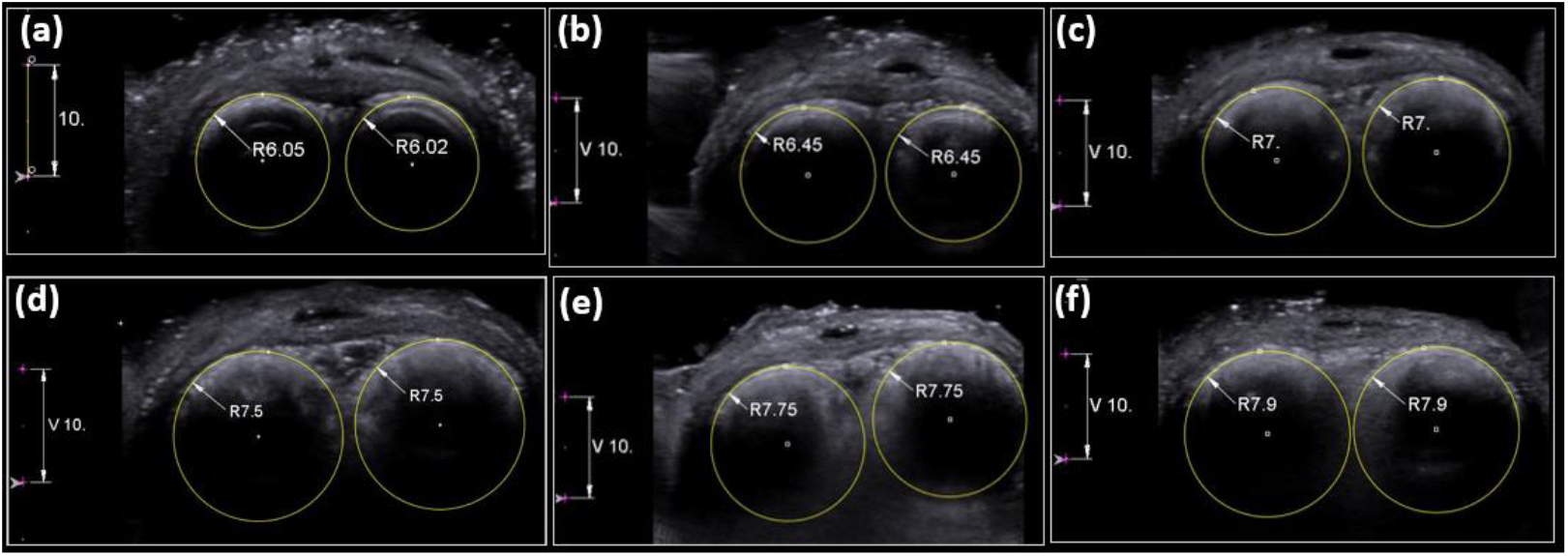
Ultrasound images and IPP cylinder diameters for distal cross-section (cross-section 1) at each inflation pressure increment, *P*: (a) *P*=0, (b) *P*=5.2 psi, (c) *P*=9.4 psi, (d) *P*=13.3 psi, (e) *P*=16.2 psi, and (f) *P*=19.1 psi. The scale bar in all images is 10mm.

**Fig. 4.**
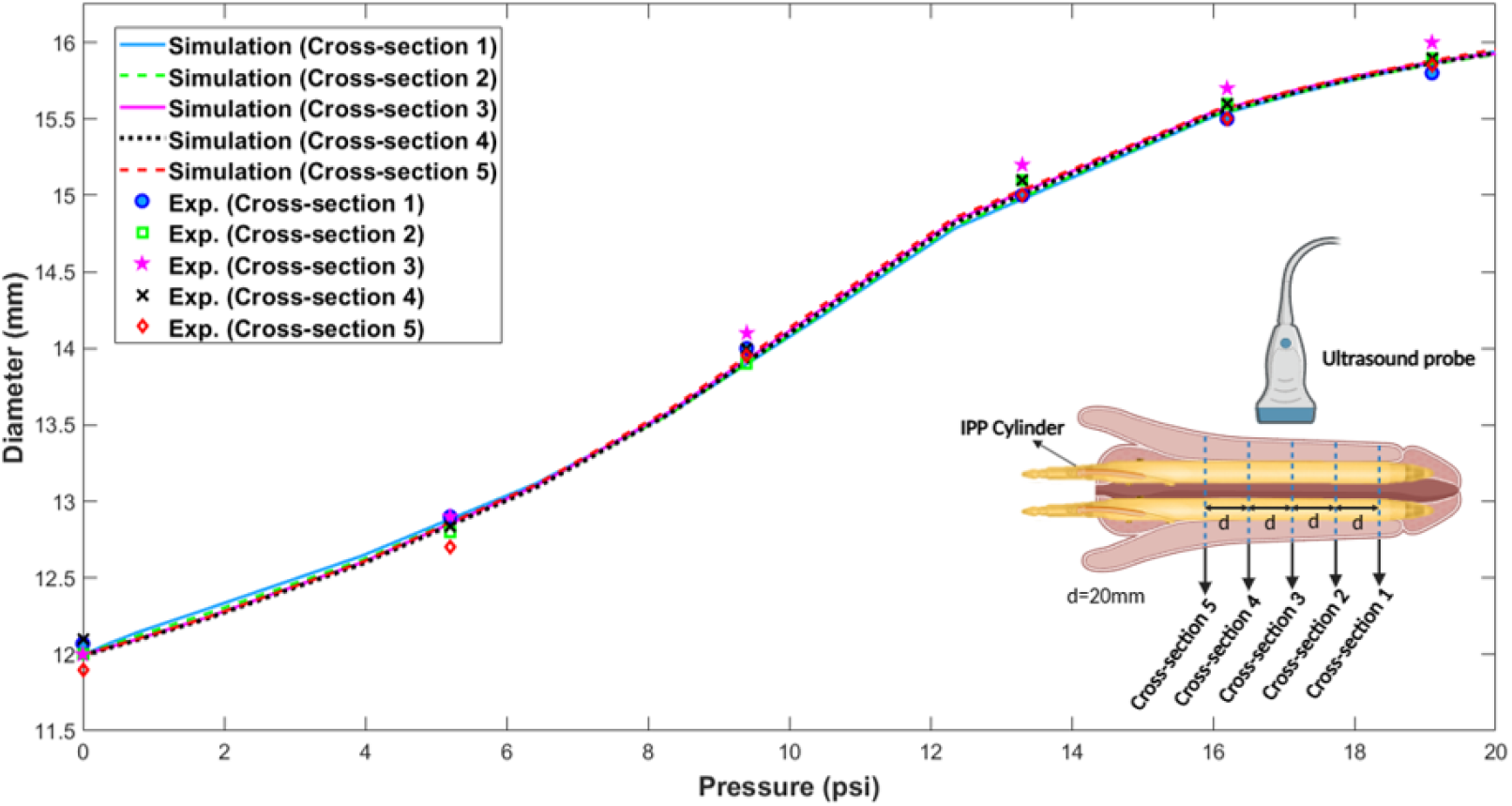
Pressure-diameter results obtained from the ex vivo experiment and in silico simulation for 5 different locations through the penile shaft.

Given this validation by the *ex vivo* experiments, this computational model can now be used to calculate and visualise device-tissue interactions, enabling observation of the mechanical behaviour of penile tissue, including displacement, stress, and strain during IPP inflation, see Fig. 5 and Fig. 6. The magnitude of tissue deformation and the loading of different penile tissue layers during IPP inflation were evaluated using displacement, maximum absolute principal logarithmic strain, and maximum absolute principal stress contours. Because the penile cross-section varies morphologically along its length, the influence of geometry on these contours was assessed by comparing a distal cross-section (without TA septum) with a proximal cross-section (with TA septum). Furthermore, two maximum inflation pressures were analysed to investigate the effect of peak pressure on the mechanical response of penile tissues. Note that the absolute maximum principal strains and stresses are used to show both maximum compressive and tensile strains at the same contour.

**Fig. 5.**
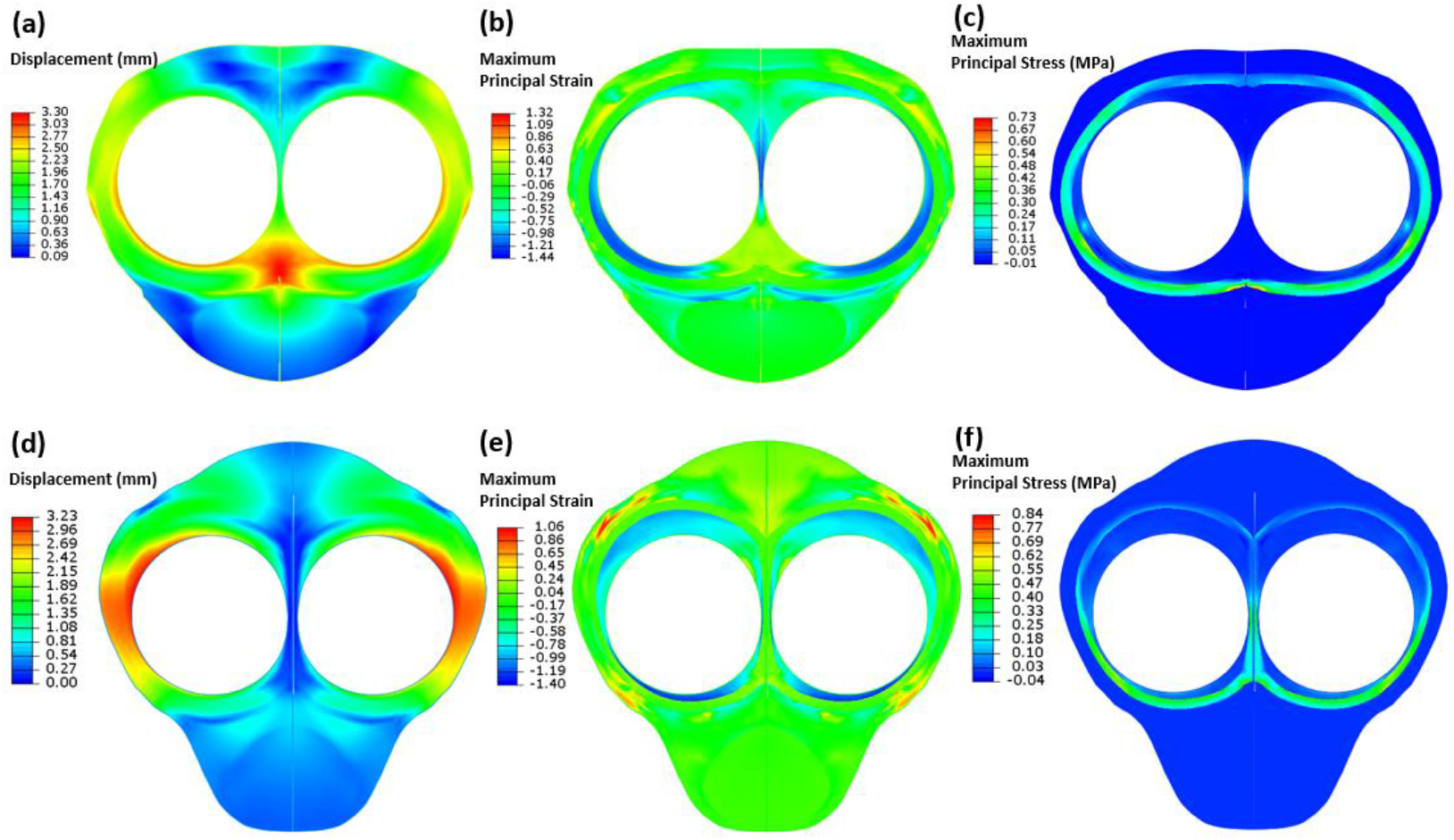
At *P*=20 psi, (a) displacement, (b) maximum absolute principal logarithmic strain, and (c) maximum absolute principal stress contours for distal cross-section (without TA septum), and (d) displacement, (e) maximum absolute principal logarithmic strain, and (f) maximum absolute principal stress contours for proximal cross-section (with TA septum).

**Fig. 6.**
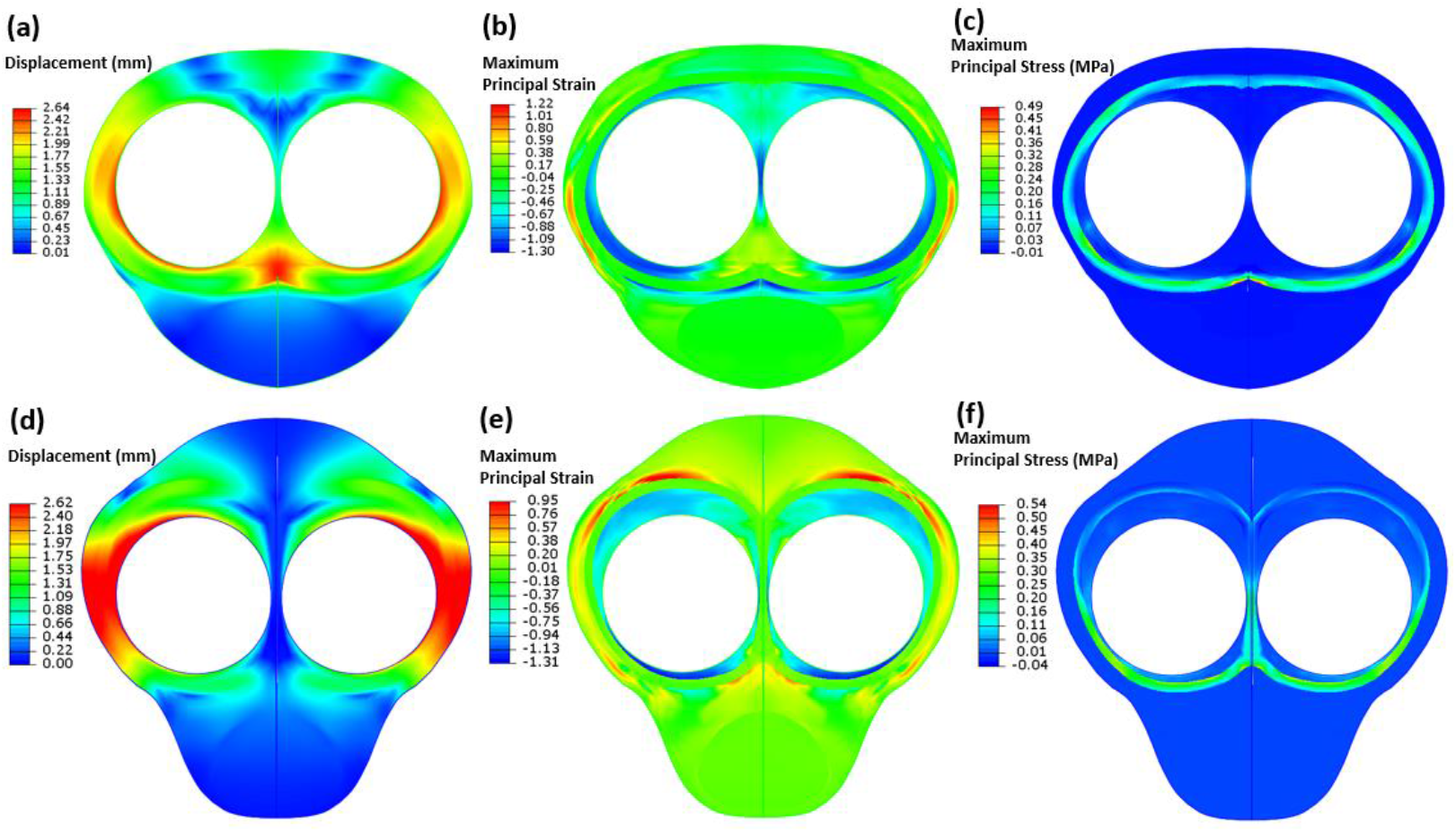
At *P*=16 psi, (a) displacement, (b) maximum absolute principal logarithmic strain, and (c) maximum absolute principal stress contours for distal cross-section (without TA septum), and (d) displacement, (e) maximum absolute principal logarithmic strain, and (f) maximum absolute principal stress contours for proximal cross-section (with TA septum).

## 4. Discussion

Comparing the results of this study with those from our previous research [28], it can be seen that the IPP cylinders inflate less when within the tissue (constrained condition) compared to inflation outside the tissue (non-constrained condition). This difference occurs because the tissue resists the cylinder’s inflation, inducing stresses and strains in the tissue, see Figs. 5 and 6. Additionally, the *ex vivo* experimental data indicates that the IPP cylinder inflation is consistent across different locations (cross-sections 1 to 5), suggesting that inflation is not region-dependent. However, the computational model reveals that tissue reactions vary slightly across different locations due to anatomical differences, as evidenced by the stress and strain contours in Figs. 5 and 6.

The computational results show that the CC layer experiences the highest compressive strain due to its spongy structure and lower stiffness compared to the cylinders and the surrounding TA layer. In contrast, the TA layer undergoes less strain but bears the highest stress, as it serves as the primary load-bearing structure due to its collagen fibre networks [29]. Consequently, as inflation pressure or diameter increases, the compression in the CC and stress in the TA layer also increase. By comparing stress and strain contours at different inflation pressures (Figs. 5 and 6), it becomes evident that inflation pressure significantly influences tissue mechanical responses. At lower pressures, 16 psi compared to 20 psi, the tissue exhibits reduced levels of stress and strain, with the maximum compressive strains in cross-sections 1 and 5 decreasing by approximately 10% and 6.5%, respectively. Similarly, the maximum principal stresses in the TA layer decrease more significantly at 16 psi, by about 32% and 35% for cross-sections 1 and 5, respectively.

The mechanical responses calculated by the computational model identify high stress/strain zones, which can help pinpoint areas vulnerable to damage during inflation. This information is valuable for informing IPP device design, allowing for the development of devices that minimise tissue damage while maintaining desired girth and stiffness.

A synthetic penile model [28] can also be developed to complement the computational model in order to investigate tissue-device interactions. Such preclinical testbeds can be improved to simulate different implantation scenarios [20], capture inelastic effects [30], compare different devices, etc. By reducing reliance on human tissues, these models help overcome ethical concerns and lower the financial burden associated with traditional experimental studies.

### 4.1. Limitations

While ultrasound imaging allows for visualisation of the device inside the tissue, it only provides a partial cross-sectional image due to the size of the penis. As a result, the full cross-section of the cylinders cannot be observed. However, it is possible to approximate the cylinder’s diameter by plotting circles from the arcs visualised in the ultrasound images. Developing an advanced technique to visualise a larger area with ultrasound would improve the accuracy of these measurements.

The computational model used in this study assumes uniform properties for each tissue layer throughout the entire penis. In cases of diseased tissues, such as Peyronie’s disease [31], [32], these properties can vary significantly. Additionally, the model assumes symmetrical geometry for each cross-section. In patients with significant anatomical differences, such as asymmetrical CC, IPP inflation would vary between the left and right corpora [20]. The model can be advanced in the future to incorporate patient-specific geometries and could also be adapted to account for region-specific tissue properties or local stiffness variations.

This study focuses on simulating IPP inflation within the penile shaft using hyperelastic models (elastic response). However, inelastic behaviours such as tissue damage and viscoelasticity are necessary for a more comprehensive analysis of device-tissue interactions. Ongoing research by the authors aims to characterise the viscoelastic and damage properties of the tissue and the IPP device, with the goal of developing a more robust computational model.

## 5. Conclusions

This study incorporates an *ex vivo* IPP implantation approach and a developed *ex vivo* IPP testing setup to establish an in silico preclinical testbed which investigates the device-tissue interactions. Comparative analysis showed that the computational model can accurately simulate the experimental IPP inflation test. This model provides valuable information of load and deformation induced in the tissue during IPP inflation. The findings highlight the ability of the model to replicate experimental behaviour and provide valuable insights into tissue mechanics that can guide device design and improve clinical outcomes.

## Acknowledgements/Funding sources

This publication has emanated from research conducted with the financial support of Science Foundation Ireland (SFI) under grant number 12/RC/2278_2 and Boston Scientific Limited (Clonmel).

